# Neurobiological Correlates of Behavioral Resilience to Chronic Alcohol and Acute Stress in Male and Female Rats

**DOI:** 10.64898/2026.01.06.697997

**Authors:** Izabela Caliman, David P. Lyvers, Jennifer Mangrum, Ayelet Kleinerman, Susan Sangha

## Abstract

**Background:** Chronic alcohol use and stress exposure are risk factors for psychiatric disorders, often leading to emotion dysregulation and impaired behavioral inhibition. While stress-related disorders are often comorbid with alcohol use disorder, not all individuals exposed to stress or alcohol develop either condition. One approach to assessing impaired behavioral inhibition of fear and reward behaviors is through conditioned inhibition paradigms.

**Methods:** Sixty-three male and seventy-five female Long Evans rats were assigned to either an intermittent two-bottle choice paradigm (alcohol versus water) or a water-only control condition, followed by exposure to an acute stressor or a non-stress control context. Subsequently, all subjects underwent a cue discrimination and conditioned inhibition learning protocol, after which brain tissue was collected for immunohistochemical analyses.

**Results:** Stress-exposed groups exhibited delayed acquisition of discrimination learning but ultimately achieved accurate differentiation among reward, fear, and inhibitor cues. When the inhibitor cue was presented concurrently with the fear cue, females subjected to both alcohol and stress failed to suppress fear responses, unlike all other groups. Notably, subsets of animals exposed to stress and/or alcohol demonstrated resilient behavioral phenotypes, which were associated with significant interregional correlations of parvalbumin and PKCδ within the corticolimbic network.

**Conclusions:** Our conditioned inhibition task for fear and reward uncovered nuanced patterns of behavioral regulation in response to fear, reward, and inhibitory cues, particularly when these cues were presented concurrently. These patterns enabled the classification of resilient versus non-resilient phenotypes, which were linked to distinct shifts in correlation profiles among interneuron subtypes.

## Introduction

Chronic alcohol consumption and stress exposure can disrupt emotion regulation and contribute to the development of psychiatric disorders. Following exposure to traumatic events, individuals may experience a range of debilitating symptoms, including intrusive recollections, avoidance behaviors, emotional numbing, and heightened arousal ^1^. Emerging research indicates these symptoms may stem from neurobiological disruptions in stimulus discrimination. When individuals struggle to distinguish between threatening and non-threatening cues, they may exhibit exaggerated responses to neutral stimuli ^2^. Stress disorders are frequently accompanied by other psychiatric conditions, with alcohol abuse or dependence occurring in approximately 41.8% of affected individuals ^3,4^. Chronic alcohol use may further compromise stimulus discrimination, potentially exacerbating symptoms related to generalization ^5^. Importantly though, not everyone exposed to these experiences develop alcohol use or stress-associated disorders ^6^.

We have developed and validated in rodents and humans an emotion regulation task that involves conditioned inhibition of fear and reward expectations ^7–14^. The rodent task starts with learning to discriminate amongst fear (paired with shock), reward (paired with sucrose) and inhibitor (paired with no outcome) cues. Then, during a test for conditioned inhibition, the inhibitor cue is co-presented with the reward cue (reward+inhibitor cue) to test for the ability of the inhibitor cue to downregulate sucrose seeking, and to test its effectiveness to downregulate fear, the inhibitor cue is also co-presented with the fear cue (fear+inhibitor cue) on separate trials. In our prior work in alcohol-free and stress-free conditions we have shown the infralimbic cortex (IL) and basolateral amygdala (BLA) contain neurons that selectively respond to the fear+inhibitor cue, indicating these regions are involved in encoding the inhibitor cue and neurons within these regions discriminate among the cues ^7,15^. Notably, we demonstrated that the infralimbic projection to the central amygdala (CeA) is necessary for the downregulated fear during the fear+inhibitor cue, indicating this specific projection is necessary for expression of cued safety ^16^.

Neural activity is heavily shaped by local interneurons, and within the medial prefrontal cortex (mPFC), parvalbumin-expressing (PV) and somatostatin-expressing (SOM) are the two of the most abundant subtypes of GABAergic interneurons ^17^. Both PV+ and SOM+ interneurons of the mPFC receive direct input from the BLA which triggers feedforward inhibition back to the BLA. Within the CeA, interneurons expressing SOM or protein kinase C-delta (PKCδ+) are predominant, reciprocally inhibiting each other to bidirectionally influence fear expression, with PKCδ+ interneurons mediating low defensive/fear states and SOM+ interneurons mediating high defensive/fear states (reviewed in ^18^). PKCδ+ interneurons of the CeA project directly to PKCδ+ interneurons of the BNST ^19^, and BNST PKCδ+ interneurons are implicated in coping with stress ^19^. Thus, the complex expression of fear and regulation of fear can be heavily influenced by local interneuron activity.

Here, we exposed male and female Long Evans rats to either a two-bottle choice of alcohol versus water or just water conditions before exposing them to an acute stress episode or control conditions ^20^. All rats went through the same discrimination learning and conditioned inhibition behavioral procedure. Voluntary alcohol or control water conditions were maintained throughout. After the test for conditioned inhibition, brains were extracted for immunohistochemical analyses of parvalbumin, somatostatin, and PKCδ interneurons within the PFC, amygdala, BNST and lateral septum. We hypothesized that behavior would be negatively impacted by alcohol and/or stress and this would be associated with changes in local interneuron networks.

## Methods and Materials

### Subjects

63 male and 75 female Long Evans rats (46-49 days old upon arrival; Envigo, Livermore, CA) were single-housed with enrichment upon arrival and allowed to acclimate for 1 week prior to any handling. Rats were then handled daily for 1 week. Estrous phases were determined on the final behavioral conditioning day. All procedures were implemented during the light cycle (07:00 to 19:00 lights on). All animal procedures were approved by the Indiana University School of Medicine Animal Care and Use Committee.

### 5 week baseline intermittent 2-bottle choice

As we have done previously ^20^, at 60-63 days of age, male rats were given 22-24g of chow per day, and females 20-22g, at the time of bottle exchange or immediately after a behavioral session. Intermittent access to both 15% alcohol and water (2-bottle choice) also started, 24h at a time, 3 times per week (bottles on Monday, Wednesday, Friday; bottles off Tuesday, Thursday, Saturday) for 5 weeks (**Figure 1A**) as we have done previously ^20^. A separate, visually distinct, water bottle was available on remaining days. Alcohol and water bottles were weighed at the end of each 24h 2-bottle choice period. The positions (left/right) of the alcohol and water bottles were randomized and all cages remained on the same housing rack along with an empty cage with alcohol and water bottles to measure and account for any spillage and/or evaporation. Rats were briefly handled and weighed at the beginning of each 24h 2-bottle choice period.

**Figure 1.**
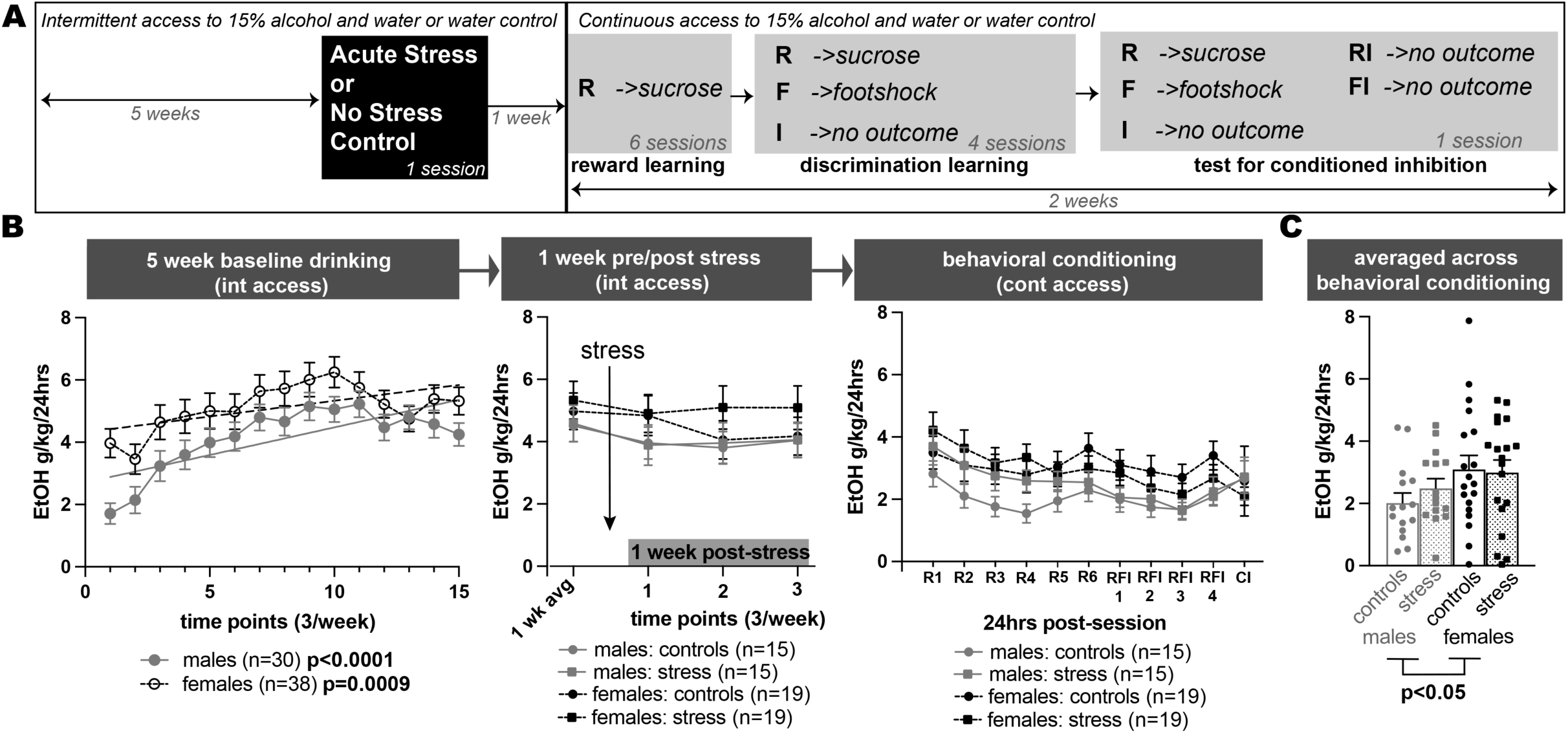
Alcohol Consumption. **A.** Schematic of experimental design of study. Rats had either intermittent access to 15% alcohol and water or just water for 5 weeks before and 1 week after a single session of either Acute Stress or No Stress Control. All groups then underwent reward learning, discrimination learning and a test for conditioned inhibition. Alcohol groups were switched to continuous access once reward learning began. **B.** Alcohol consumed (g/kg/24hrs) across 1) the 5-week baseline, 2) 1 week after stress exposure, and 3) across behavioral conditioning. **C.** Averaged alcohol consumed (g/kg/24hrs) across all of behavioral conditioning.

### Stress exposure

After 5 weeks of home cage intermittent 2-bottle choice, rats were exposed to either Stress or No Stress conditions (**Figure 1A**). As we have done previously ^20,21^, stress consisted of a single 90min session in Context A in which 15 unsignaled 1s, 1mA footshocks were presented (variable ITI, 4-8mins). No Stress conditions consisted of a single 90min session in Context A without any footshocks presented. Standard MedAssociate operant conditioning chambers served as Context A with background houselights off and a cotton ball doused with vanilla extract placed within the sound attenuating chamber but outside of the Plexiglas box to introduce an odor to the context. Footshocks were delivered through a grid floor via a constant current aversive stimulator. This yielded 4 groups per sex: control (i.e. water conditions with no stress), stress alone, alcohol alone, and alcohol+stress. For the 1 week following Stress or No Stress exposure, rats were returned to the same intermittent 2-bottle choice schedule and procedure as described above (**Figure 1A**).

### Conditioned inhibition task

One week after stress or control exposure, all rats completed a conditioned inhibition task in Context B (**Figure 1A**). Context B used the same MedAssociate boxes but with houselights on and no vanilla odor. Sessions were video-recorded for offline scoring. Three cues were used: a 20s white noise (reward), a 20s pulsing 11kHz tone (fear), and two 20s cue lights (inhibitor). Stimuli were not counterbalanced, but our prior work shows no learning differences among these stimuli ^7,11^. At the start of conditioning (R1), rats switched to continuous 2-bottle choice (alcohol/water), with daily weighing of bottles and rats; consumption data reflect the 24h period after each session. Rats received six reward sessions (R1–R6), each with 20 reward cue–sucrose pairings (ITI 90–130s; sucrose delivered 10–18s after cue onset). During R6, rats were pre-exposed to five trials each of fear and inhibitor cues only to assess baseline freezing without inducing latent inhibition ^7–11,21,22^. One day later, rats underwent four reward–fear–inhibitor sessions (RFI1–4), each with 15 reward trials (sucrose at 18s), four fear trials (0.5s, 0.5mA footshock at cue offset), and 20 inhibitor trials (no shock) (39 trials total, ITI 100–140s). Finally, one day after RFI4, rats completed a test for conditioned inhibition with 28 trials (pseudorandom presentation, ITI 100-140s): reward (6 trials with sucrose), fear (4 trials with footshock), inhibitor (6 trials), fear+inhibitor (6 trials, no shock), and reward+inhibitor (6 trials, no sucrose).

### Immunohistochemistry

Rats were anesthetized 90mins after the final session with isoflurane and transcardially perfused with phosphate buffered solution (PBS) and 4% paraformaldehyde in 0.1M PBS. Brain tissue preparation, slicing (35µm coronal sections), immunohistochemistry, and immunofluorescence procedures were carried out as described previously ^23,24^. We performed staining for PV, SOM, and PKCδ, using the following primary and secondary antibodies: Mouse Anti-Parvalbumin (SAB4200545, Sigma-Aldrich, USA, 1:30000), Mouse Anti-Somatostatin (GTX71935, GeneTex, USA, 1:500), Rabbit Anti-PKCδ (ab182126, Abcam, Germany, 1:750), biotinylated goat anti-rabbit, and horse anti-mouse antibody (BA-1000-1.5 and BA-2000-1.5, respectively, Vector Laboratories, USA, 1:500).

Photomicrographs were obtained with a Leica microscope (DM6B, Leica microsystems, USA) with a 20x objective. Specific brain structures analyzed included (mm AP relative to bregma): PFC (PL, IL, DP, and Cg1): +3.24 to +3.0, BNST and lateral septum: +0.12 to -0.12, and amygdala (CeA, LA, and BLA): -2.28 to -2.52. Immunopositive cells were counted either manually by a person blind to the experimental procedures or automatically through ImageJ (version 1.54p) or Qupath (version 0.5.1) software. For data presented as mean fluorescence intensity, the following formula was used: Corrected total cell fluorescence = Integrated Density – (area of selected brain structure x mean fluorescence of background readings). For each brain region, blind analyses were conducted bilaterally.

### Analyses

All statistical analyses and graphs were carried out and generated using GraphPad Prism 10.6.1 (GraphPad Software Inc., San Diego, CA). and are presented as the mean ± SEM.

#### Ethanol consumption

Ethanol and water intake were measured over 24 hours, with spilled amounts subtracted from consumption. Alcohol intake was expressed as grams of ethanol per kilogram of body weight per 24-hour session (ETOH g/kg/24h), accounting for ethanol density. Data were analyzed using linear regression, two- and three-way repeated measures ANOVAs, followed by LSD post hoc tests where appropriate.

#### Behavior

Behaviors were scored manually from video recordings using custom Python scripts that masked cue and animal IDs to ensure blind scoring. Fear was measured as freezing, defined as complete immobility except for respiration ^25,26^, and expressed as the percentage of time spent freezing during each cue. Sucrose seeking was assessed by the percentage of time spent at or inside the reward port during each cue. Behavioral data were analyzed using three-way repeated measures ANOVAs with Dunnett’s post hoc tests where appropriate.

#### Resilient/Non-resilient phenotypes

Freezing and port-seeking behaviors during conditioned inhibition were used to classify alcohol- and/or stress-exposed subjects as resilient or non-resilient. Fear and reward suppression ratios were calculated as the percentage of time freezing or port-seeking to the cue paired with an inhibitor divided by that to the cue alone (i.e. FI/F or RI/R). Lower ratios indicated stronger inhibition, while ratios above 0.8 reflected persistent fear or reward-seeking despite the inhibitor cue. Subjects with FI/F > 0.8 (‘persistent fear’, PF), RI/R > 0.8 (‘persistent reward’, PR), or both (PFR) were classified as non-resilient.

#### Immunohistochemistry

Data are presented as number of positive neurons/mm^2^ for all analyses, except PKCδ and SOM expression in CeA, where data are presented as corrected total cell fluorescence. Data were analyzed using linear regression and two-way ANOVA, followed by Tukey’s post hoc tests where appropriate.

## Results

### Alcohol consumption

Prior to stress or behavioral conditioning, male (M; n=30) and female (F; n=38) rats significantly increased alcohol consumption over the 5-week baseline period of intermittent access to two-bottle choice (simple linear regression, *M: p<0.0001, F: p<0.001;* **Figure 1B**). Drinking behavior did not significantly change in response to the Stress or No Stress experience for the 1 week following compared to 1 week prior (**Figure 1B**). When subjects were moved to continuous access throughout behavioral conditioning the Stress and Control groups did not differ from each other within sex (**Figure 1B**), however, there was a main effect of sex with females consuming more alcohol than males (*2-way ANOVA, F(1,64)=4.2, p=0.04*) (**Figure 1C**).

### Behavioral Conditioning

#### Reward-Fear-Inhibitor Discrimination Learning

One week after the Stress/No Stress exposure rats underwent 6 sessions of reward conditioning followed by four sessions of reward-fear-inhibitor discrimination learning (**Figure 1A**), where the reward cue was paired with sucrose, the fear cue with footshock, and the inhibitor cue with no outcome. Groups were Control (M n=16, F n=18), Stress (M n=17, F n=19), Alcohol (M n=15, F n=19), and Alcohol+Stress (M n=15, F n=19).

Freezing during each cue was expressed as a percentage. Three-way ANOVAs were run for each sex per session (**Table 1**). In the first discrimination session (RFI1), males showed significant alcohol × stress × cue and stress × cue interactions, plus a main effect of cue. Females showed a stress × cue interaction and main effects of stress and cue. Post hoc Dunnett’s tests revealed that non-stressed males and females froze significantly more to the fear cue than to the inhibitor or reward cues (p < 0.01; **Figure 2A**). In contrast, stressed animals (with or without alcohol) did not discriminate fear in this session. In the last discrimination learning session (RFI4), males and females showed a significant stress x cue interaction and a main effect of cue. Post hoc Dunnett’s to the fear cue showed all groups froze significantly more to the fear cue than all other cues (*p-values <0.05*) (**Figure 2B**).

**Figure 2.**
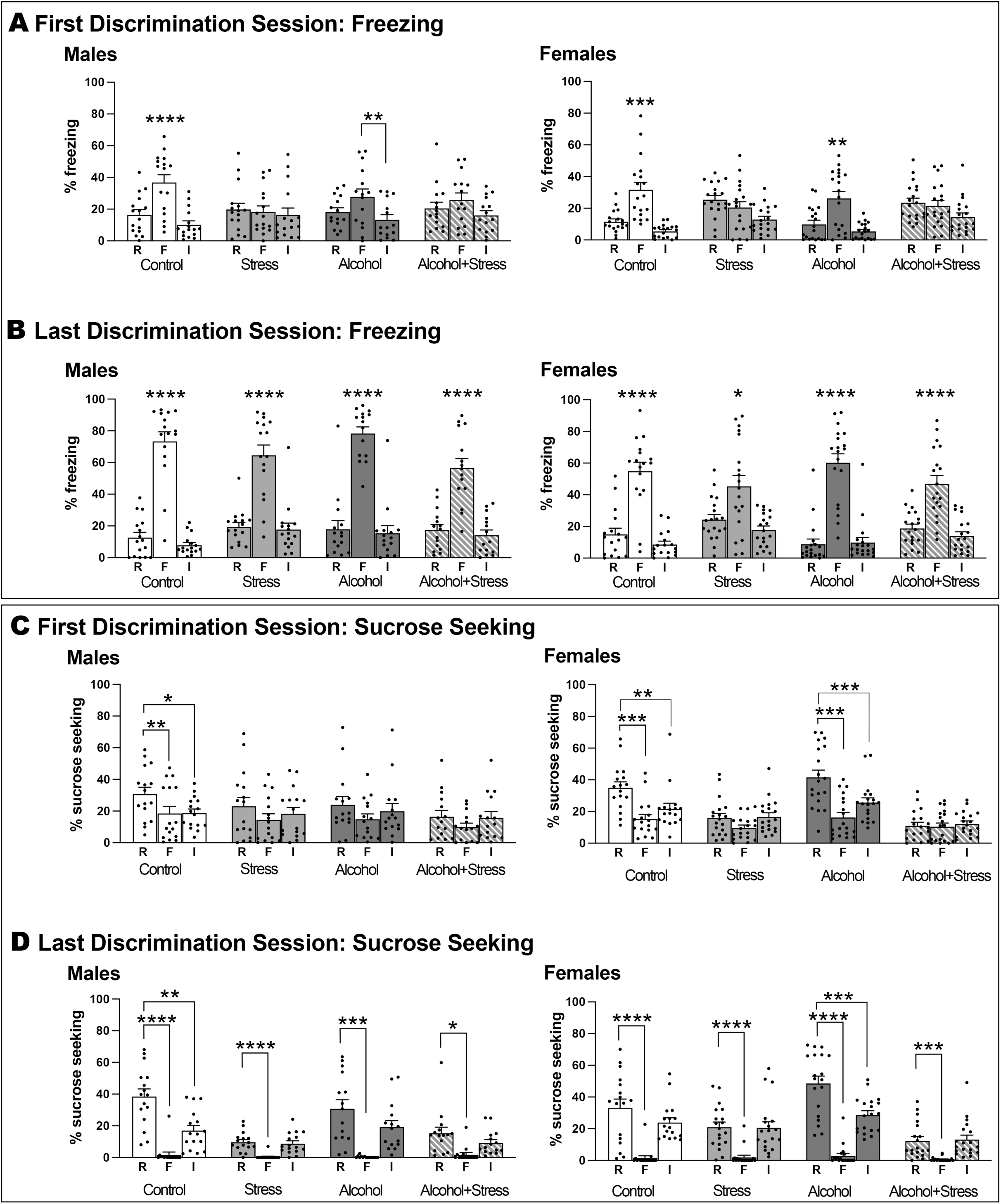
Stress delayed discrimination learning. Percent time freezing during each cue for the first (**A**) and last (**B**) cue discrimination sessions are shown for males (left) and females (right) across control, stress, alcohol and alcohol+stress conditions. *p<0.05, **p<0.01, ***p<0.001, ****p<0.0001 compared to F cue. Percent time sucrose seeking during each cue for the first (**C**) and last (**D**) cue discrimination sessions are shown for males (left) and females (right) across control, stress, alcohol and alcohol+stress conditions. *p<0.05, **p<0.01, ***p<0.001, ****p<0.0001 compared to R cue.

**Table 1.** Summary of significant effects on discrimination learning.

| **Table 1. RFI Discrimination Learning (Figure 2)** |  |  |
| --- | --- | --- |
| **First Session: Freezing** |  |  |
| Males,<br>3-way ANOVA | alcohol x stress x cue | F(2,116)=4.58, p=0.01 |
|  | stress x cue | F(2,116)=9.9, p=0.0001 |
|  | ME cue | F(1.5, 85.2)=27.27, p<0.0001 |
|  | post hoc Dunnett's to F cue | Control: F>R, p<0.0001; F>I, p<0.0001<br>Alcohol: F>I, p<0.01 |
| Females,<br>3-way ANOVA | stress x cue | F(2,142)=20.72, p<0.0001 |
|  | ME stress | F(1,71)=5.57, p=0.02 |
|  | ME cue | F(1.3,92.1)=39.2, p<0.0001 |
|  | post hoc Dunnett's to F cue | Control: F>R, p<0.001; F>I, p<0.001<br>Alcohol: F>R, p<0.01; F>I, p<0.01 |
| **Last Session: Freezing** |  |  |
| Males,<br>3-way ANOVA | stress x cue | F(2,116)=8.19, p=0.0005 |
|  | ME cue | F(1.2,70.8)=254.6, p<0.0001 |
|  | post hoc Dunnett's to F cue | Control: F>all cues, p<0.0001<br>Stress: F>all cues, p<0.0001<br>Alcohol: F>all cues, p<0.0001<br>Alcohol+Stress: F>all cues, p<0.0001 |
| Females,<br>3-way ANOVA | stress x cue | F(2,140)=9.34, p=0.0002 |
|  | ME cue | F(1.1,80.5)=133, p<0.0001 |
|  | post hoc Dunnett's to F cue | Control: F>all cues, p<0.0001<br>Stress: F>all cues, p<0.05<br>Alcohol: F>all cues, p<0.0001<br>Alcohol+Stress: F>all cues, p<0.0001 |
| **First Session: Sucrose Seeking** |  |  |
| Males,<br>3-way ANOVA | ME cue | F(1.6,90)=12.05, p<0.0001 |
|  | post hoc Dunnett's to R cue | Control, R>F, p<0.01; R>I, p<0.05 |
| Females,<br>3-way ANOVA | stress x cue | F(1.8,124)=18.44, p<0.0001 |
|  | ME stress | F(1,70)=41.73, p<0.0001 |
|  | ME cue | F(1.8,124)=30.37, p<0.0001 |
|  | post hoc Dunnett's to R cue | Control: R>F, p<0.001; R>I, p<0.01<br>Alcohol: R>F, p<0.001; R>I, p<0.001 |
| **Last Session: Sucrose Seeking** |  |  |
| Males,<br>3-way ANOVA | stress x cue | F(2,116)=17.01, p<0.0001 |
|  | ME stress | F(1,58)=29.65, p<0.0001 |
|  | ME cue | F(1.6,94.5)=69.64, p<0.0001 |
|  | post hoc Dunnett's to R cue | Control: R>F, p<0.0001; R>I, p<0.01<br>Stress: R>F, p<0.0001<br>Alcohol: R>F, p<0.001<br>Alcohol+Stress: R>F, p<0.05 |
| Females,<br>3-way ANOVA | alcohol x stress x cue | F(2,140)=4.25, p=0.02 |
|  | alcohol x stress | F(1,70)=9.91, p=0.002 |
|  | stress x cue | F(2,140)=21.04, p<0.0001 |
|  | ME stress | F(1,70)=31.45, p<0.0001 |
|  | ME cue | F(1.9,130)=119, p<0.0001 |
|  | post hoc Dunnett's to R cue | Control: R>F, p<0.0001<br>Stress: R>F, p<0.0001<br>Alcohol: R>F, p<0.0001; R>I, p<0.001<br>Alcohol+Stress: R>F, p<0.001 |

Sucrose seeking was measured as time at the port during each cue (% of cue). Three-way ANOVAs were run for each sex per session (**Table 1**). In RFI1, males showed a main effect of cue, while females showed a stress × cue interaction and main effects of stress and cue. Stress reduced sucrose seeking in females. Post hoc tests revealed Control males and females and Alcohol females sought sucrose significantly more during the reward cue than other cues (**Figure 2C**). In contrast, all stress groups and Alcohol males failed to show reward discrimination in this session. In RFI4, males showed a stress × cue interaction and main effects of stress and cue. Females exhibited interactions of alcohol × stress × cue, alcohol × stress, and stress × cue, along with main effects of stress and cue. Post hoc tests indicated all groups sought sucrose more during the reward cue than the fear cue, with Control males and Alcohol females also showing higher sucrose seeking than during the inhibitor cue (*p-values <0.05*) (**Figure 2D**).

#### Test for Conditioned Inhibition of Fear and Reward Seeking

All male and female groups showed significant inhibition of freezing during the fear+inhibitor cue compared to the fear cue (*p-values <0.05*), with the exception of the female Alcohol+Stress group who showed similar levels of freezing to the fear cue and fear+inhibitor cue (**Figure 3A**). Three-way ANOVAs performed (**Table 2**) for each sex for freezing behavior showed a significant stress by cue interaction in the females and main effects of cue for both sexes. Three-way ANOVAs performed for each sex for sucrose seeking showed both sexes had significant stress by cue interactions and main effects of stress and cue. In addition, the males also showed significant alcohol by stress and alcohol by stress by cue interactions. Overall, stress groups showed dampened sucrose seeking compared to non-stress groups (**Figure 3B**).

**Figure 3.**
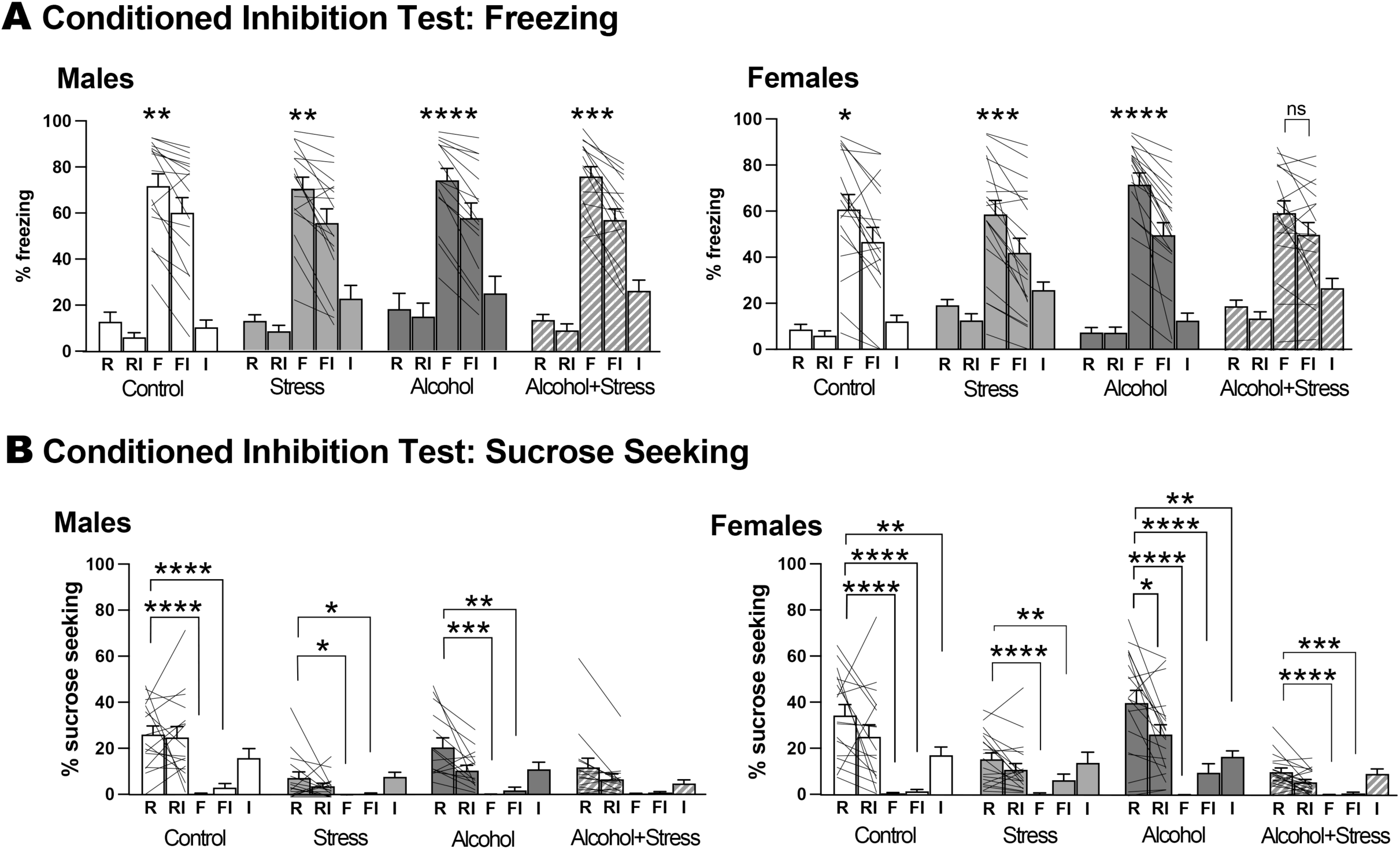
Females exposed to alcohol and stress did not show conditioned Inhibition of fear. **A.** Percent time freezing during each cue shown for males (left) and females (right) across control, stress, alcohol and alcohol+stress conditions. *p<0.05, **p<0.01, ***p<0.001, ****p<0.0001 compared to F cue. **B.** Percent time sucrose seeking during each cue shown for males (left) and females (right) across control, stress, alcohol and alcohol+stress conditions. *p<0.05, **p<0.01, ***p<0.001, ****p<0.0001 compared to R cue.

**Table 2.**
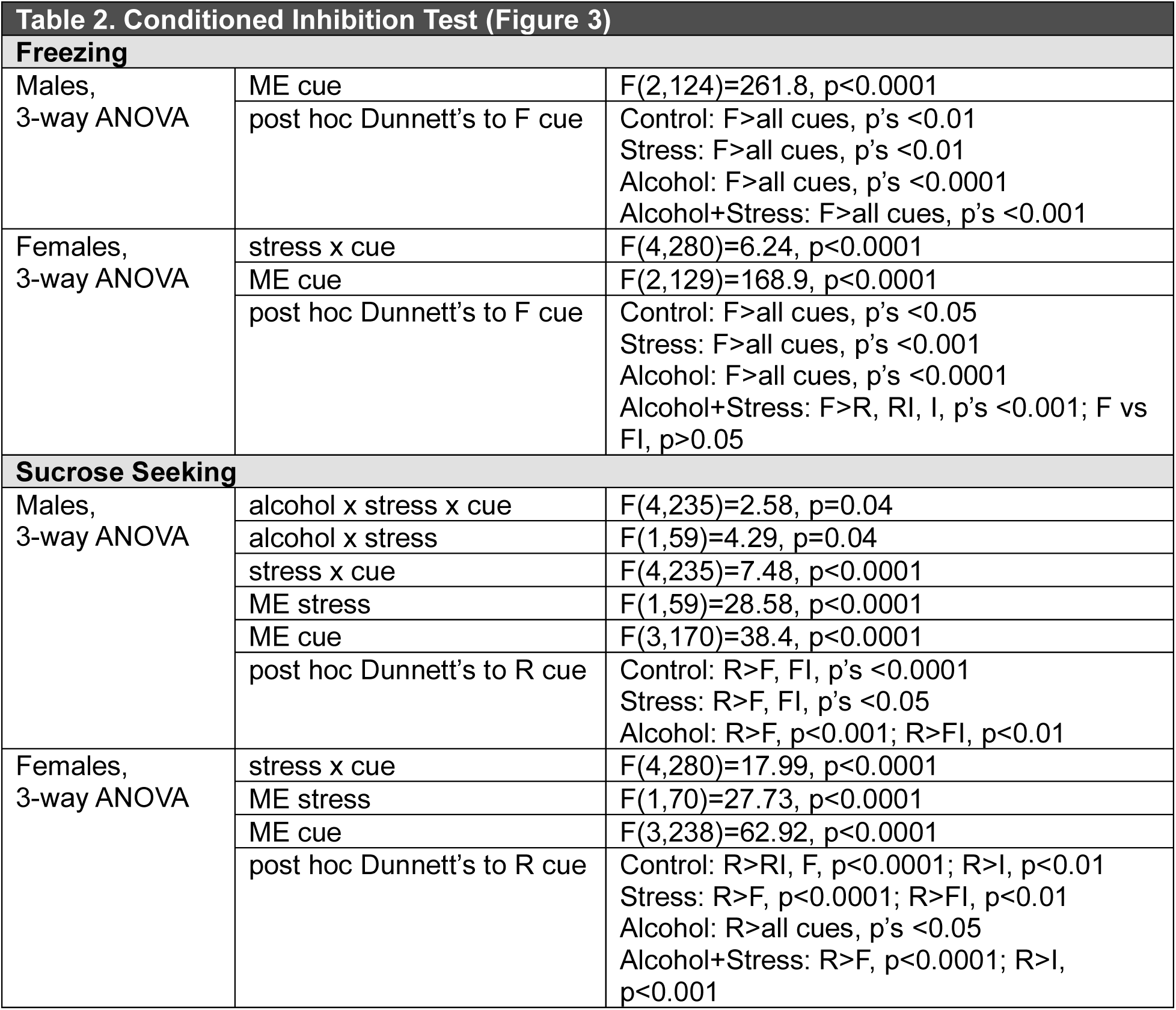
Summary of significant effects on conditioned inhibition.

### Immunohistochemistry

Brains were extracted and 35µm coronal sections of the medial PFC (Cg, PL, IL, DP), amygdala (LA, BA, CeA), BNST and lateral septum were stained for one of the following: parvalbumin (PV), somatostatin (SOM), or PKCδ.

For each subregion of the mPFC, we observed main effects of alcohol in males in number of PV+ interneurons (**Figure 4A-D**). Two-way ANOVAs for each sex was performed for the effects of stress and alcohol on number of PV+ interneurons for the Cg, PL, IL and DP (**Figure 4A-D**; **Table 3)**. There were significant stress by alcohol interactions for PL, IL, and DP, as well as main effects of alcohol in males for Cg, PL, IL and DP. For each region, alcohol conditions showed lower number of PV+ interneurons than non-alcohol conditions in males. This was also negatively correlated with alcohol consumption in males for each subregion (**Figure 4A-D**). That is, alcohol consumption correlated negatively with the number of PV+ interneurons for Cg (*p=0.01*), PL (*p=0.009*), IL (*p=0.006*), and DP (*p=0.03*). None of these measures were significant in females (**Figure 4A-D**). For the same subregions of the mPFC, the number of SOM+ interneurons (**Figure 4E-H**; **Table 3**) showed no significant changes with the exception of main effects of alcohol in the male PL and IL, and a negative correlation between the number of SOM+ interneurons in the DP with alcohol consumption in males (*p=0.03*). None of these measures were significant in females. There were also no significant differences in number of PV+ or SOM+ neurons in the lateral amygdala, basal amygdala or central amygdala, with the exception of a main effect of alcohol on SOM+ neurons in the lateral amygdala of females **(Supplemental Figures 1-2; Table 3).**

**Figure 4.**
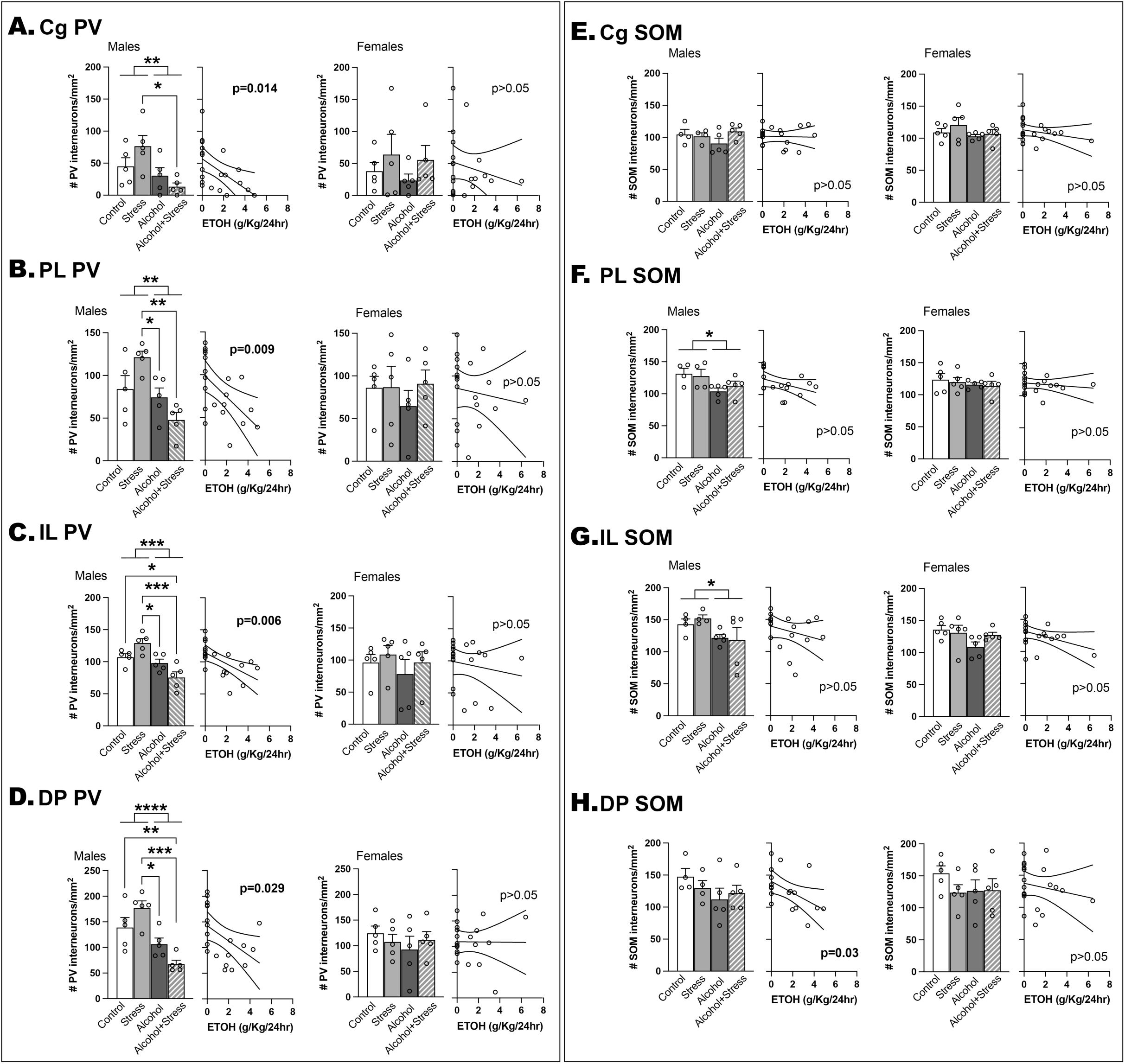
Males exposed to alcohol showed reduced number of PV+ cells in the prefrontal cortex that was correlated with alcohol consumption. Number of PV+ interneurons are shown for Cg (**A**), PL (**B**), IL (**C**) and DP (**D**) for males and females across control, stress, alcohol and alcohol+stress conditions (left) and by amount of alcohol consumed (right). Number of SOM+ interneurons are shown for Cg (**E)**, PL (**F**), IL (**G**) and DP (**H**) for males and females across control, stress, alcohol and alcohol+stress conditions (left) and by amount of alcohol consumed (right). *p<0.05, **p<0.01, ***p<0.001, ****p<0.0001.

**Table 3.**
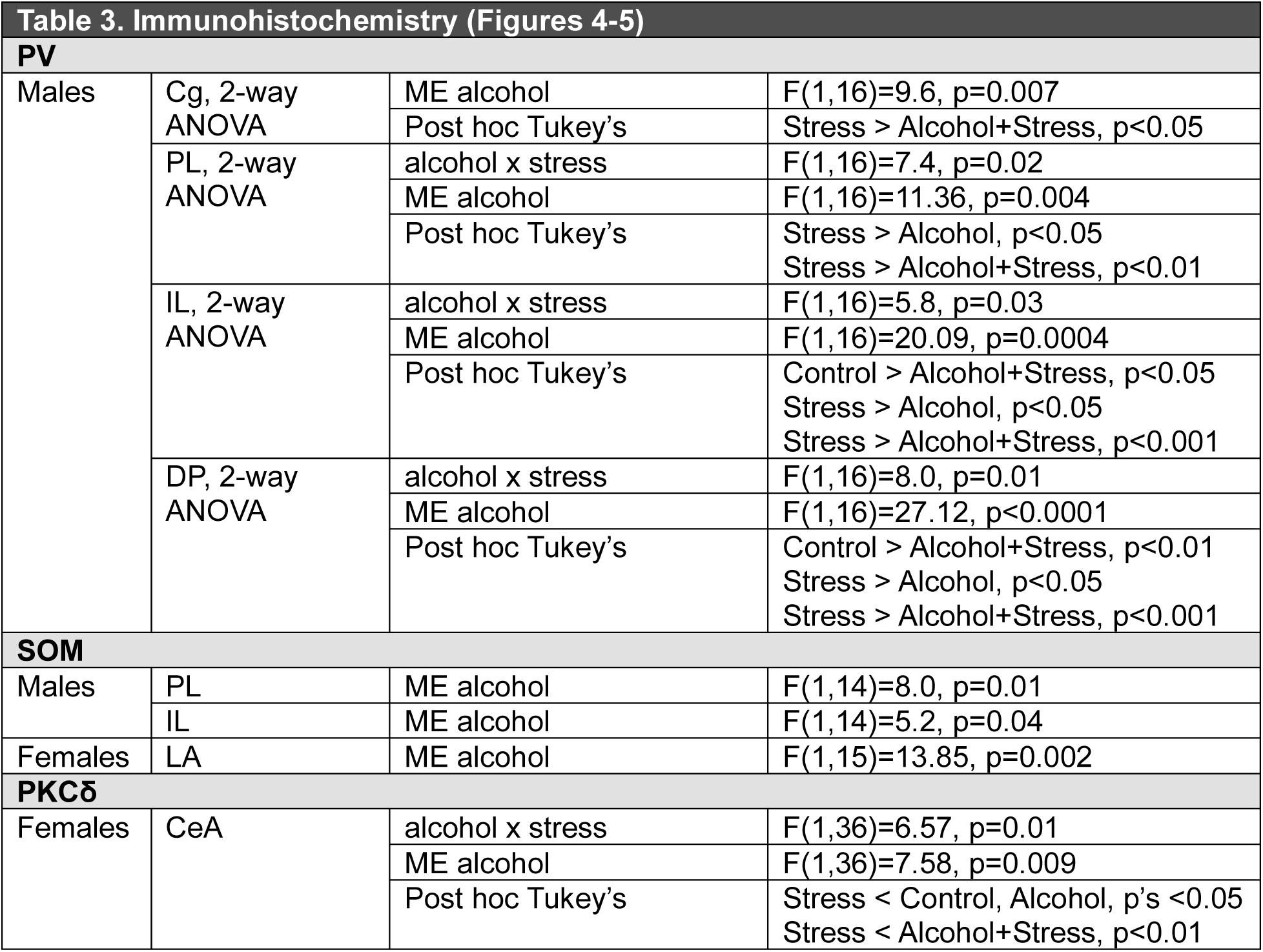
Summary of significant effects on PV, SOM and PKCδ in prefrontal cortex and amygdala.

Two-way ANOVAs for each sex was performed for the effects of stress and alcohol on PKCδ interneuron expression for the CeA, BNST and lateral septum (**Figure 5**; **Table 3**). Within the female CeA there was a significant stress by alcohol interaction and main effect of alcohol. Post hoc Tukey’s showed PKCδ was significantly reduced in females exposed to stress compared to control (*p=0.02*), alcohol alone (*p=0.01*), and alcohol+stress (*p=0.003*) groups. These changes were not correlated with alcohol consumption. Similar analyses for PKCδ in the female BNST or lateral septum showed no significant effects, as well as no significant differences in the male CeA, BNST or lateral septum.

**Figure 5.**
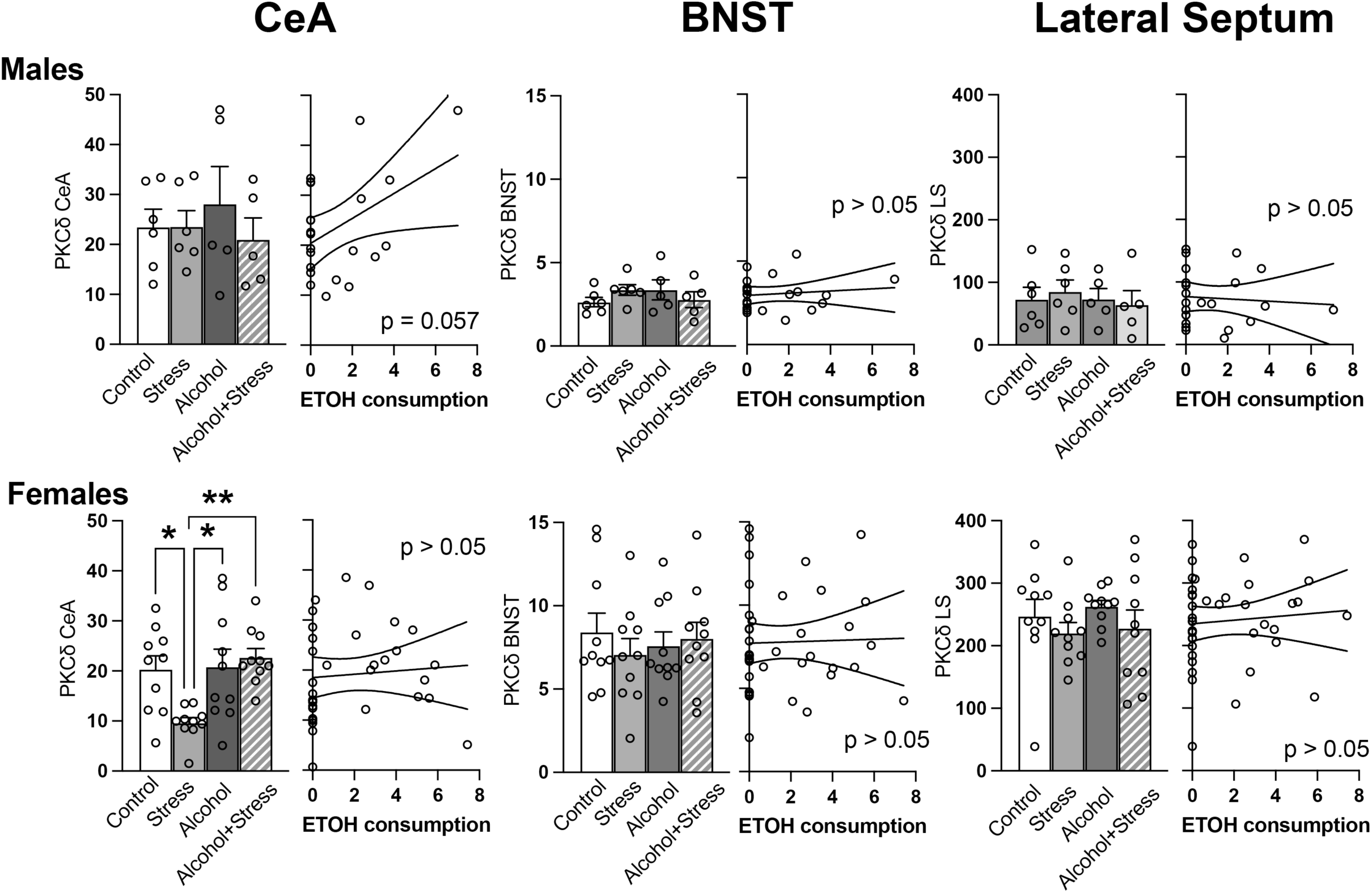
Females exposed to stress showed reduced PKCδ in the central amygdala. Corrected total cell fluorescence for PKCδ are shown for CeA, BNST and lateral septum for males and females across control, stress, alcohol and alcohol+stress conditions (left) and by amount of alcohol consumed (right). *p<0.05, **p<0.01.

### Resilient/Non-resilient Phenotypes

#### Behavior of resilient versus non-resilient phenotypes

Freezing and port-seeking during conditioned inhibition were used to classify subjects exposed to alcohol and/or stress as resilient or non-resilient (**Figure 6**). Suppression ratios were calculated as FI/F (‘fear+inhibitor’/’fear’) and RI/R (‘reward+inhibitor’/’reward’); lower ratios indicated stronger inhibition.

**Figure 6.**
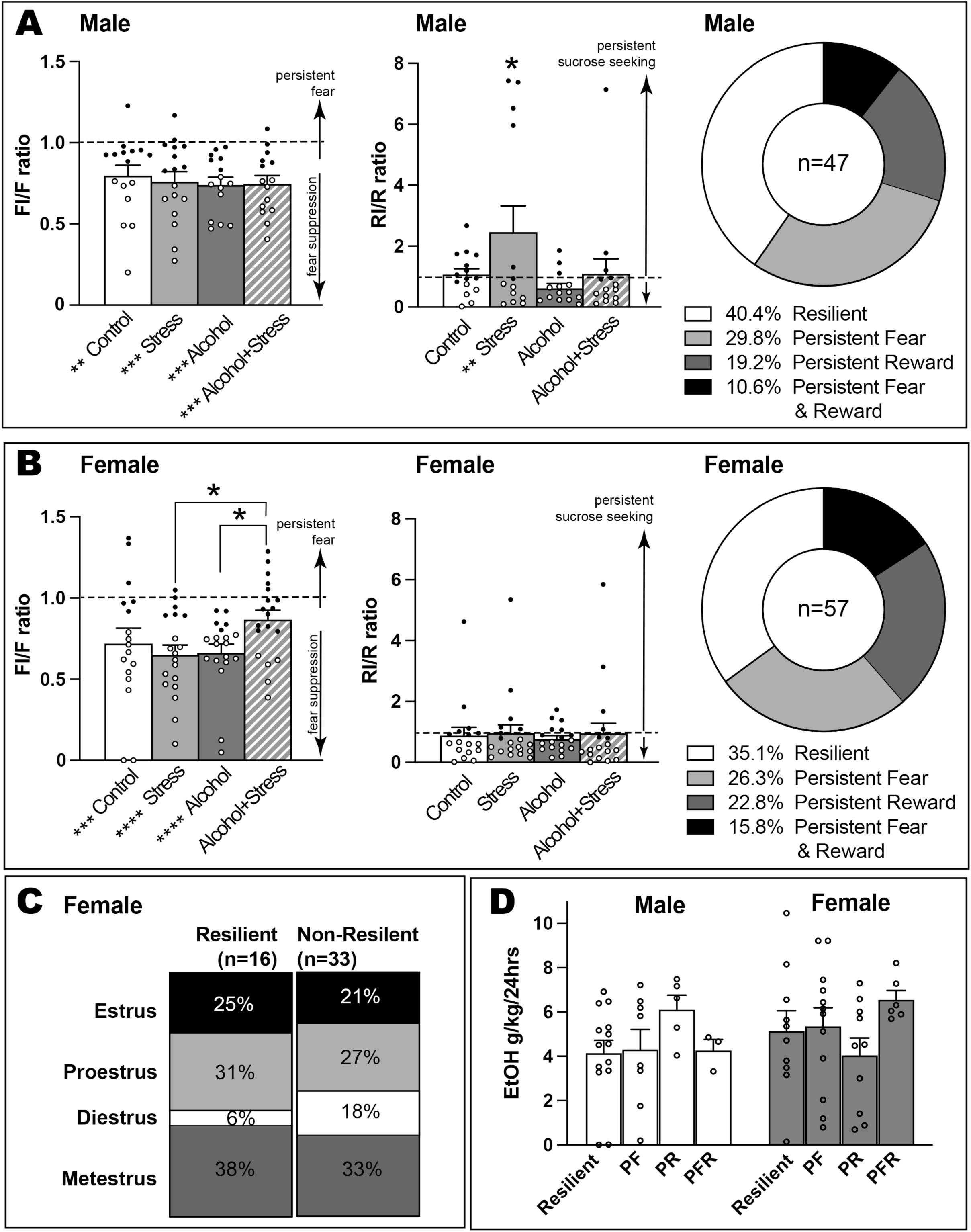
Conditioned inhibition behaviors as resilient vs non-resilient phenotypes. Conditioned inhibition of fear and reward ratios shown across control, stress, alcohol and alcohol+stress conditions. Ratios <0.8 were classified as non-resilient: persistent fear, persistent reward or both. **A.** In males, FI/F ratios for all groups were significantly lower than 1.0, while the RI/R ratios for the stress group were significantly higher than 1.0 and all other groups (*p<0.05). Across all males exposed to alcohol and/or stress (n=47), 40.4% showed ratios <0.8 and classified as resilient. **B.** In females, the only group that did not show FI/F ratios being significantly lower than 1.0 was the alcohol+stress group, which was also significantly higher than the stress and alcohol groups (*p<0.05). Across all females exposed to alcohol and/or stress (n=57), 35.1% showed ratios <0.8 and classified as resilient. **C.** Proportion of females in diestrus on the day of testing was higher in the non-resilient subjects (persistent fear, persistent reward or both). **D.** Baseline alcohol consumption prior to stress was not significant across phenotypes but males classified as persistent reward and females classified as persistent fear and reward had elevated baseline alcohol consumption. PF, persistent fear; PR, persistent reward; PFR, persistent fear and reward. In **A** and **B:** black circles represent non-resilient subjects (suppression ratios >0.8); * in the X axis (group conditions) represents significantly different from 1.0.

For fear inhibition, FI/F ratios in males (**Figure 6A, left**) were similar across conditions (main effect of alcohol *F(2,90) = 19.32, p<0.0001*), all significantly below 1.0 (*p’s <0.002*). In females, (**Figure 6B, left)**, there was an alcohol x stress interaction (*F(2,106)=3.62, p=0.03*) and main effect of alcohol (*F(2,106)=18.82, p<0.0001*). Alcohol+stress females had higher FI/F ratios than stress alone (*p=0.004*) and alcohol alone (*p = 0.007*) and was the only group not different from 1.0 (*p=0.08*), unlike control (*p=0.0004*), stress alone (*p<0.0001*), and alcohol alone (*p<0.0001*) groups.

For reward inhibition (**Figure 6A, middle**), male stress-alone rats showed elevated RI/R (*p<0.01* vs all groups; *p=0.006* vs 1.0), with main effects of alcohol (F(2,83)=3.41, p=0.04) and stress (*F(1,83)=4.3, p=0.04*), indicating persistent reward seeking to the reward cue despite the presence of the inhibitor. Females showed no significant differences (**Figure 6B, middle)**.

Subjects with suppression ratios >0.8 were classified as non-resilient by displaying persistent fear and/or reward seeking despite the inhibitor cue being present (black circles in **Figure 6A-B, left and middle**). Among alcohol and/or stress-exposed rats, 40.4% of males and 35.1% of females were resilient; more females showed “persistent fear and reward” than males (15.8% vs 10.6%) (**Figure 6A-B, right**). Diestrus females were more often non-resilient on the day of conditioned inhibition testing (18% vs 6%) (**Figure 6C**). Baseline drinking one week prior to stress/no-stress exposure and subsequent behavioral conditioning did not differ significantly by phenotype, although it did appear males classified as ‘persistent reward’ (PR) and females classified as ‘persistent fear and reward’ (PFR) may have had slightly higher baseline alcohol consumption (**Figure 6D**).

#### Immunohistochemistry of resilient versus non-resilient phenotypes

We re-analyzed immunohistochemistry by resiliency phenotype and tested if PV, SOM, and PKCδ levels were correlated across PFC and amygdala networks (**Figure 7**). All significant correlations reported below were in the positive direction. **PV correlations (Figure 7A):** Resilient males (n=9) showed strong interregional PV correlations (Cg, PL, IL, DP; p<0.05, r>0.66 except Cg–IL), whereas non-resilient males (n=5) had fewer correlations (PL–Cg p<0.01, r=0.97; PL–IL p<0.05, r=0.89; IL–DP p<0.05, r=0.90). Resilient females mirrored non-resilient males (PL–Cg p<0.01, r=0.89; PL–IL p<0.01, r=0.85; IL–DP p<0.05, r=0.83). Non-resilient females (n=7) showed no significant PV correlations. **SOM correlations (Figure 7B):** Minimal interregional correlations were observed; resilient males showed Cg–DP correlation (p<0.05, r=0.70), while non-resilient females showed PL–Cg (p<0.05, r=0.61) and PL–IL (p<0.05, r=0.85) correlations. **PKCδ correlations (Figure 7C):** Resilient males (n=7) and females (n=8) exhibited BNST–CeA correlations (p<0.01, r=0.91, 0.87), which were absent in non-resilient subjects. Non-resilient females instead showed BNST–LS correlations (combined n=22, p<0.01, r=0.63; PF subgroup n=11, p<0.05, r=0.63; PFR subgroup n=7, p<0.05, r=0.77).

**Figure 7.**
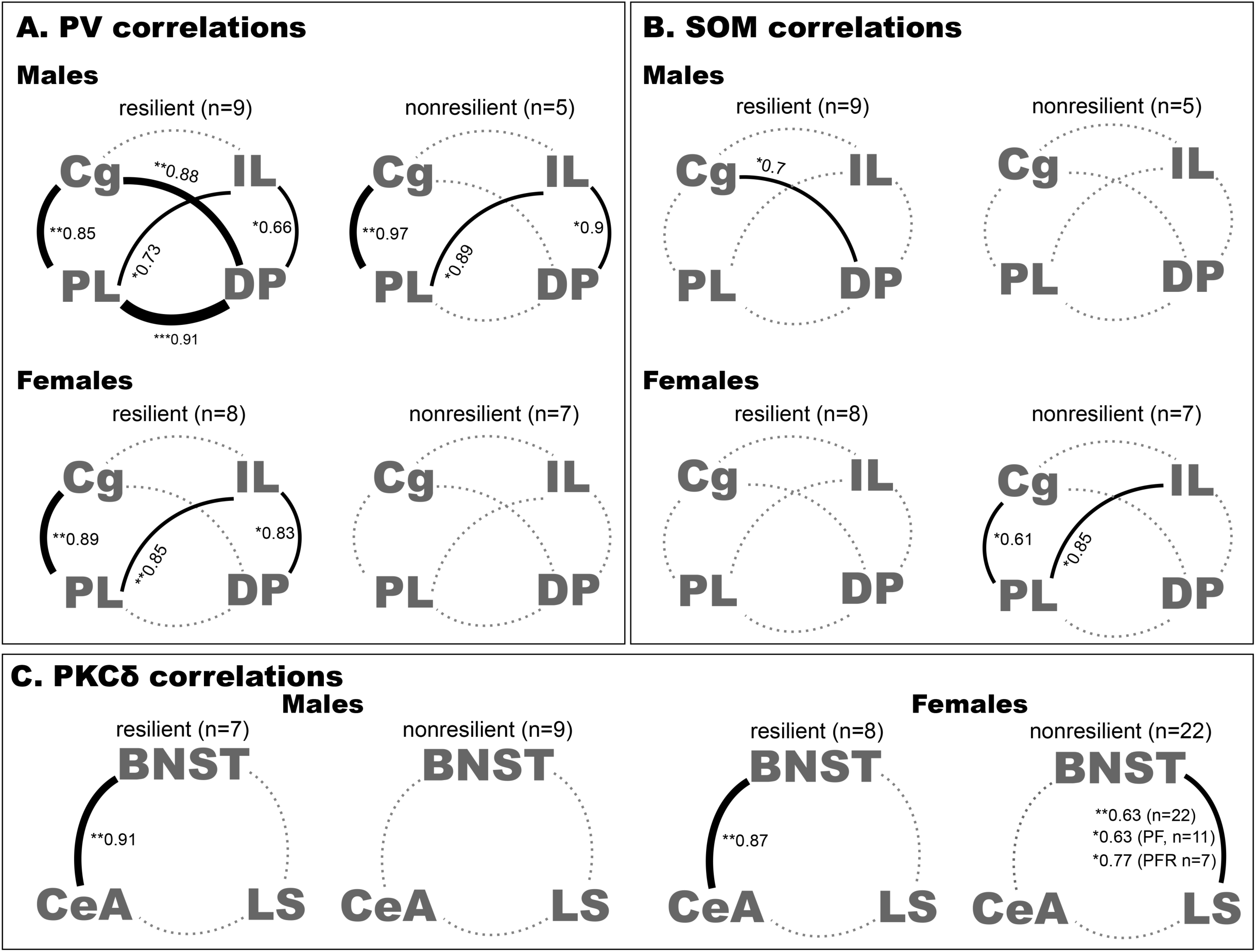
Resiliency phenotype was associated with PV, SOM, and PKCδ correlations across PFC and amygdala networks. Significant correlations between brain regions are shown as r-values (*p<0.05, **p<0;01, ***p<0.001) and solid lines. The weight of the lines reflects the level of significance. Dashed lines indicate correlations were not significant. **A.** PV correlations showed resilient males (n=9) had strong interregional PV correlations, with the exception of Cg-IL, whereas non-resilient males (n=5) had fewer correlations. Resilient females mirrored non-resilient males and non-resilient females (n=7) showed no significant PV correlations. **B.** SOM correlations from the same subjects as **A** showed minimal interregional correlations. Resilient males showed a Cg–DP correlation, while non-resilient females showed PL–Cg and PL–IL correlations. **C.** PKCδ correlations showed resilient males (n=7) and females (n=8) exhibited BNST–CeA correlations, which were absent in non-resilient subjects. Non-resilient females instead showed BNST–LS correlations (all non-resilient subjects combined n=22, **p<0.01, r=0.63; PF subgroup n=11, *p<0.05, r=0.63; PFR subgroup n=7, *p<0.05, r=0.77).

## Discussion

Our findings demonstrate that chronic alcohol exposure and acute stress interact to produce distinct behavioral and neurobiological outcomes. Behaviorally, stress impaired early discrimination between fear, reward, and inhibitor cues, while alcohol further modulated these effects. Our prior work similarly showed stress exposure reduced sucrose seeking and delayed cue discrimination ^20^. Although most groups eventually learned to discriminate cues, stress consistently dampened sucrose-seeking behavior, and females exposed to both alcohol and stress uniquely failed to inhibit fear responses when the inhibitor cue was present. That is, all male and female groups showed significant inhibition of freezing during the fear+inhibitor cue compared to the fear cue, with the exception of the female Alcohol+Stress group (**Figure 3A**). This points to a sex-specific vulnerability in fear regulation under combined alcohol and stress conditions. Neurobiologically, alcohol reduced parvalbumin-positive interneuron counts across medial prefrontal cortex subregions in males, with reductions correlating negatively with recent alcohol consumption. In comparison, somatostatin-positive interneurons showed minimal changes, while PKCδ-positive interneurons in the central amygdala were significantly reduced in stressed females, independent of alcohol intake. These findings highlight differential circuit-level adaptations to alcohol and stress across sexes.

When we took into account individual differences in behavioral outcomes, resilience analyses revealed that 40% of males and 35% of females maintained effective fear and reward inhibition despite prior alcohol and/or stress exposure. Non-resilient phenotypes were more prevalent among females in diestrus, suggesting hormonal influences on stress-alcohol interactions. This is consistent with other studies in rodents linking anxiety-like behaviors with diestrus and metestrus (lower ovarian hormones) than proestrus ^27^. In contrast, our prior work found no relationship of estrous phase with either conditioned inhibition of fear or reward ^12^, although our prior work did not classify individual subjects based on how they inhibited both their fear and sucrose seeking behaviors. The female alcohol+stress group was the only one to not show conditioned inhibition of fear. This is largely consistent with our prior work demonstrating females may have a different threshold than males for expressing conditioned inhibition of fear^12^.

Network analyses revealed distinct patterns associated with resilience versus vulnerability. Resilient animals showed strong PV interneuron correlations across prefrontal regions and robust BNST–CeA PKCδ correlations. In contrast, non-resilient females exhibited BNST–lateral septum correlations, suggesting fundamentally different corticolimbic network configurations. Within the CeA, PKCδ+ interneurons are linked to low defensive/fear states, whereas SOM+ interneurons mediate high defensive/fear states (reviewed in ^18^). In our study, stress significantly reduced PKCδ+ interneurons in females, likely biasing the network toward heightened defensive states. Notably, PKCδ+ interneurons in the CeA project directly to PKCδ+ interneurons in the BNST ^19^, and BNST PKCδ+ interneurons have been implicated in stress coping ^19^. Thus, our finding that resilient males and females exhibit increased PKCδ+ correlations between the CeA and BNST supports the role of this subnetwork in adaptive stress regulation. Differential contributions of PL, IL, and DP to fear regulation are well established, with PL and DP generally promoting fear and IL promoting safety ^9,15,16,28,29^. Maintaining balanced output across these regions through precise coordination by local PV interneurons likely fosters resilience, consistent with our observation that resilient animals display higher PV correlations across Cg, PL, IL, and DP.

It is possible that conditioned inhibition behaviors prior to exposure to alcohol or stress are directly associated with later resilience. Our conditioned inhibition behaviors were assessed after several weeks of homecage alcohol consumption and exposure to an acute stressor. Acheson and colleagues ^30^ have shown in humans that poor discrimination between a fear CS+ and CS-prior to military deployment (i.e. stress) was associated with increased reported Post-Traumatic Stress Disorder (PTSD) symptoms after deployment. This study did not directly assess conditioned inhibition (i.e. presenting the CS+ and CS-concurrently) but their results suggest that impaired fear regulation in a discrimination paradigm is not produced by PTSD per se but perhaps is a behavioral marker of who may be susceptible to later developing PTSD symptoms in response to a stressor. While others have investigated vulnerability and resilience to developing PTSD or alcohol use disorder ^30–33^, relatively few have looked at resilient behaviors associated with fear/reward regulation to cues despite having a prior experience of stress and/or alcohol conditions.

Overall, these results underscore that conditioned inhibition tasks provide a powerful framework for identifying behavioral resilience and its underlying neural substrates. The observed sex-specific patterns suggest that resilience to alcohol and stress is mediated by unique circuit-level mechanisms, offering novel targets for interventions aimed at mitigating risk for stress- and alcohol-related psychopathology.

## Acknowledgments

This work was supported by NIMH R01MH110425 to SS. The authors would like to thank Dr. A. Oblak for access to a Leica microscope (DM6B, Leica microsystems, USA) for generating images of immunohistochemical data, and Z. Ahmed for assistance in processing tissue for immunohistochemistry. DPL, JM performed experiments. IC processed tissue for immunohistochemistry. DPL, IC, AK, SS analyzed results. SS designed the study and wrote the manuscript.

---

**Supplemental Figure 1.**
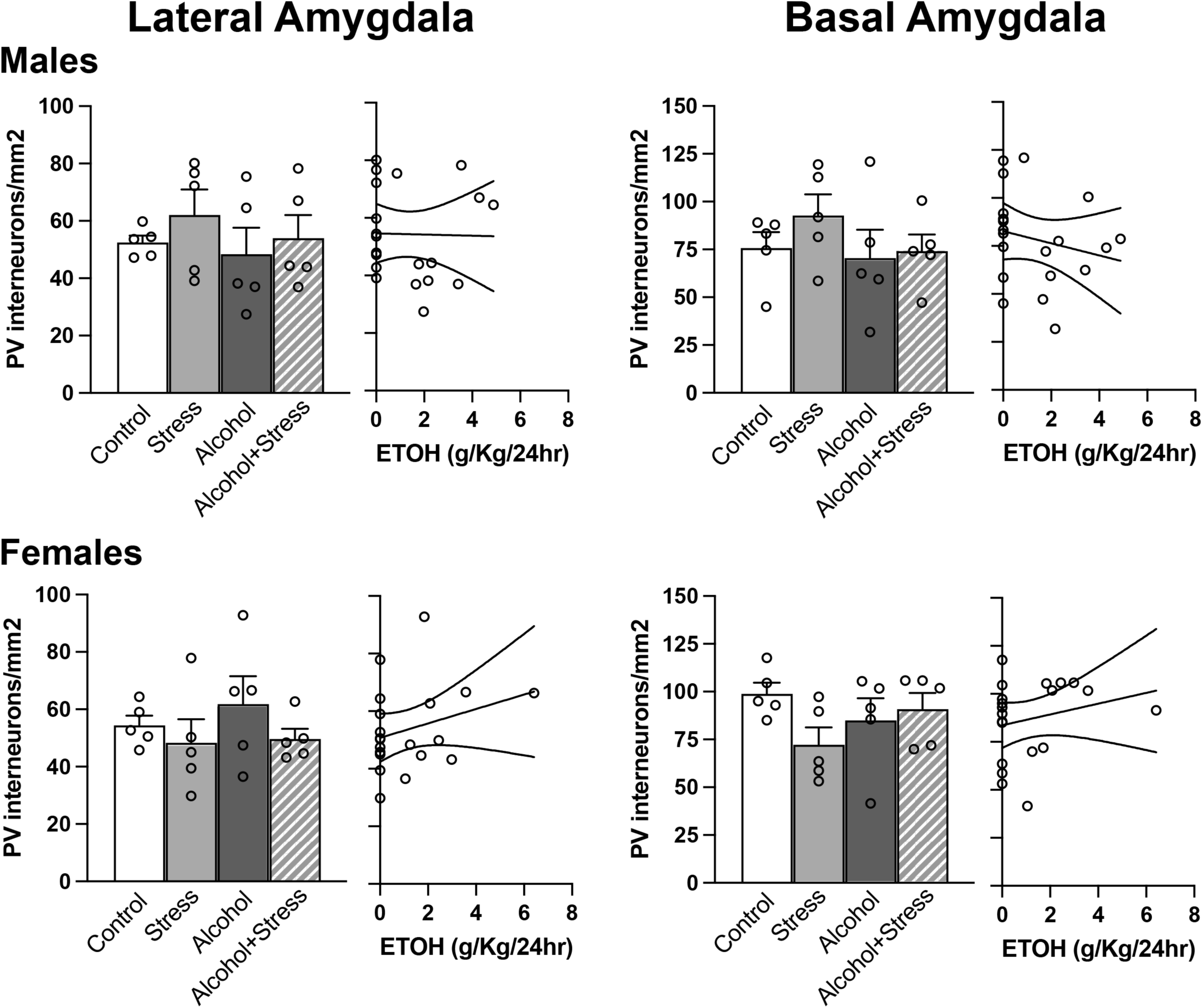
Number of PV+ cells in the amygdala. Number of PV+ interneurons are shown for lateral amygdala and basal amygdala for males and females across control, stress, alcohol and alcohol+stress conditions (left) and by amount of alcohol consumed (right). There were no significant differences.

**Supplemental Figure 2.**
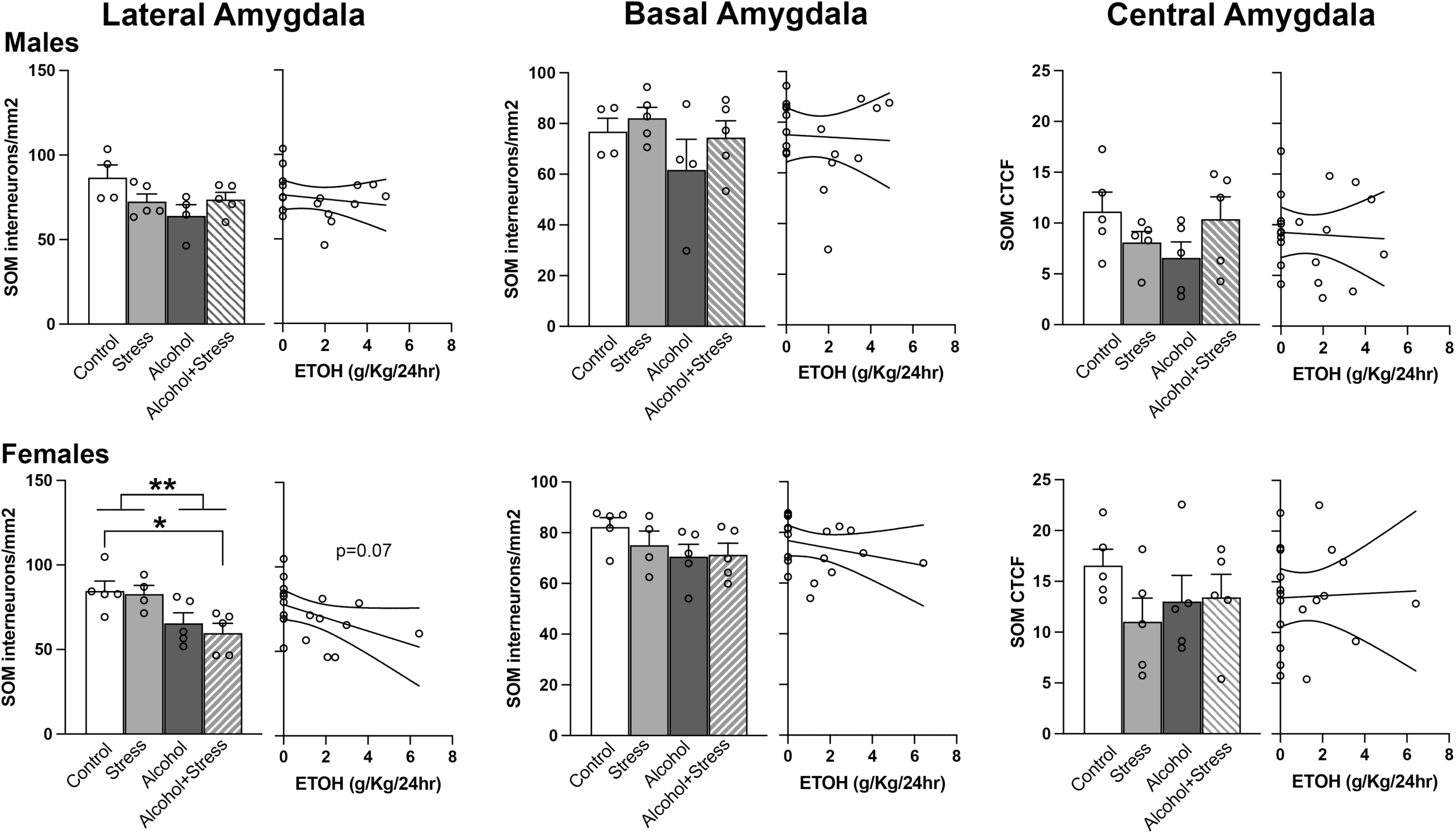
Number of SOM+ cells in the amygdala. Number of SOM+ interneurons are shown for lateral, basal, and central amygdala for males and females across control, stress, alcohol and alcohol+stress conditions (left) and by amount of alcohol consumed (right). Overall, there were no significant differences, with the exception of a main effect of alcohol in the female lateral amygdala (**p<0.01) where post hoc Tukey’s also showed alcohol+stress was significantly lower than controls (*p<0.05). CTCF, corrected total cell fluorescence.

